# Update and Expansion of the Distribution of the Maldonado Redbelly Toad, *Melanophryniscus moreirae* Gallardo 1966, in Southeast Brazil, Using Citizen Science Data

**DOI:** 10.64898/2026.01.24.701131

**Authors:** Lucas Aosf, Micheli D.M. Intir, Tayane M. de Azevedo, Leandro B.C. Menezes, Matheus T. Moroti

## Abstract

*Melanophryniscus moreirae* is a diurnal species endemic to the high-altitude grasslands of the Serra da Mantiqueira mountain range, currently classified as “Near Threatened” (NT) by the IUCN. Knowledge about its distribution and natural history is fundamental for conservation plans, especially in the face of the threats of climate change. This study presents a new record of the species in the state of Sao Paulo, at an altitude of 1,904 m, expanding the known distribution in the southwestern portion of the Serra da Mantiqueira. In addition to fieldwork, a spatial and temporal review of data available on the GBIF (Global Biodiversity Information Facility) platform was carried out. The temporal analysis confirmed observation peaks in November, coinciding with the reproductive period, and an absence of records in the colder months, consistent with the species’ dormancy behavior. The study demonstrates that the integration of citizen science data, when properly validated, is an effective tool to fill knowledge gaps about biodiversity and assist in the monitoring of threatened species.

---

*Melanophryniscus* Gallardo 1966, is a monophyletic genus divided into three clades based on their morphological characteristics: *M. moreirae* group (Miranda-Ribeiro 1920), *M. tumifrons* group (Boulenger 1905), and *M. stelzneri* group (Weyenbergh 1875) (Kwet et al. 2005; Aguirre et al. 2021). Currently, 31 species are recognized, distributed in Argentina, Uruguay, Paraguay, Bolivia, and Brazil, all within South America (Kwet et al. 2005; Van Sluys and Guido-Castro 2011; Zank et al. 2014; Aguirre et al. 2021; Frost 2025). The species of *Melanophryniscus* occupy a variety of habitats at different elevations, such as open areas, wetlands, and forests. However, most species predominantly inhabit high elevations, reaching up to 2,500 m (Guix et al. 1998; Bornschein et al. 2015), with only *M. setiba* Peloso, Faivovich, Grant, Gasparini, and Haddad 2012, known from lowland areas, specifically from a locality in the Restinga of Espírito Santo State, southeastern Brazil (Peloso et al. 2012).

*Melanophryniscus moreirae* is a diurnal species (Van Sluys and Guido-Castro 2011; Haddad et al. 2013) that inhabits high-elevation grasslands along the Serra da Mantiqueira in southeastern Brazil (Bokermann 1967; Guix et al. 1998; Carvajalino-Fernández et al. 2013), covering the States of Rio de Janeiro (Miranda-Ribeiro 1920), Minas Gerais (Weber et al. 2007; Ortiz et al. 2017), and São Paulo (Marques et al. 2006; Zank et al. 2014). Currently, the species is classified as Near Threatened (NT) by the International Union for Conservation of Nature (IUCN 2023). The population trend is currently stable, with no significant continuous decline observed in the extent or quality of its habitat. However, climate change may alter the dynamics of the high-elevation habitats where the species lives, potentially classifying it as Vulnerable (VU) in the near future. Therefore, further studies on threats, distribution, and population trends of this species are recommended (IUCN 2023).

The integration of data from various sources and citizen science platforms, such as iNaturalist data through the Global Biodiversity Information Facility (GBIF), is increasingly relevant for adding new information about species, especially regarding geographic distribution (Hochmair et al. 2020; Hewitt et al. 2021; Salvador et al. 2021). These platforms have been aiding researchers in obtaining additional data, particularly in areas that are difficult to access or have restrictions (Mesaglio et al. 2021). Moreover, this approach has already enabled the documentation of new (Winterton 2020), rare (Salvador and Cavallari 2020; Wilson et al. 2020), and even exotic species (Vendetti et al. 2018; Gladstone et al. 2020; Balashov and Markova 2021; Hausdorf et al. 2021). Additionally, the integration of these data can fill gaps in the natural history of species, provided that the available data on these platforms meet established quality standards and can be reviewed by experts (Kosmala et al. 2016).

In this context, we add a new record of *M. moreirae* for the state of São Paulo and update the known distribution of the species. Additionally, we spatially and temporally explore the data available on the GBIF platform to enhance our understanding of the species.

During a three-day expedition for wildlife observation, we recorded an individual of *M. moreirae* in the highlands of the Gomeral neighborhood at 1400 h on 3 January 2024, in the municipality of Guaratinguetá, State of São Paulo, southeastern Brazil (22º40’ S, 45º23’ W, WGS 84, elev. 1,904 m). In this location, high-elevation grassland is the predominant vegetation type. It was raining during the day, and we found the individual in a temporary pool along with several newly metamorphosed *Rhinella* sp., near rocks and typical high-elevation grassland vegetation with many grasses. A voucher specimen was collected (ICMBio provided collecting permits 87096-4) and is deposited in the Amphibian Collection of the Museum of Biological Diversity at UNICAMP (ZUEC-AMP 26551). In addition to the new record and literature data, we incorporated GBIF records to update the species distribution map. We also analyzed occurrence patterns concerning altitude, phenology, and the historical period of documentation. These parameters, acknowledged as crucial for conservation planning (Mesaglio & Callaghan, 2021), can significantly enrich our knowledge of the species’ natural history and distribution (Mesaglio and Callaghan 2021).

We extracted data from GBIF using the species name with the “*rgbif*” package in R (R Core Team 2021), reviewed and classified the obtained records into four categories: (1) “validated,” which included records with coordinates; (2) “centroid,” those that represented the centers of municipalities instead of potential occurrence areas of the species. Since the species does not occur in urban areas, these are more likely to be coordinate entry errors and were not represented on the map (except for Campos do Jordão (SP), see details below); (3) “without coordinates,” which were records without coordinates that could not be confirmed precisely for the occurrence location; and (4) “invalidated,” which were those questioned in the literature (e.g., Langone 2005) and/or with potential errors in the platform databases. For example, a disjunct population of *M. moreirae* was supposedly found in Óbidos, Pará, Brazil, and designated as *Atelopus moreirae massarti* (Cochran 1948), a subspecies that was later considered a synonym of *M. moreirae*. However, this designation was questioned due to possible confusion in the collection labels from the Massart expedition (Bokermann 1967; Guix et al. 1998; Frost 2025). Historical records indicate that the expedition also visited Itatiaia at an elevation of 2,200 m, suggesting a possible mix-up of specimens collected on the plateau and in the Pará Amazon (Langone 2005). Following Bokermann’s (1967) recommendation, Langone (2005) analyzed two paratypes of *Atelopus moreirae massarti* described by Cochran (1948) and concluded that, despite their preservation state, these specimens did not present significant morphological differences compared to those from the type locality.

Caramaschi and Pombal (2025) reviewed the type specimens of *Melanophryniscus moreirae*, designating a lectotype and defining the paralectotypes, as well as correctly georeferencing the type locality. In the same study, the authors also reassessed the taxonomic status of *Atelopus moreirae massarti*, confirming that it is a junior synonym of *Melanophryniscus moreira*e and, therefore, should not be recognized as a valid species or subspecies. Consequently, records from the dubious locality were excluded (see details in the supplementary material). The supposed type locality of *A. moreirae massarti* in the state of Pará, Brazil, was considered an error resulting from mislabeling of specimens collected during the Massart Expedition (Caramaschi and Pombal 2025).

All points were manually verified, and their elevation was recorded using the original collection notebook annotations when available (see details in supplementary material) or extracted using Google Earth®. To represent the temporal series of records, we excluded only those categorized as “invalidated” or those without dates, totaling 171 records. We created the figures using the *ggplot2* package in R (R Core Team 2021), and the map was produced using QGIS software (QGIS Development Team 2022).

We found 339 records of *M. moreirae* in the scientific literature and GBIF data, with only six of them (∼1.7 %) documented in traditional peer-reviewed scientific literature. Of the 333 GBIF records, 190 were without coordinates (∼58 %), 78 were considered validated (21.5 %), 57 records corresponded to municipality centroids (∼18 %), and eight records were considered invalidated (2.5 %). The vast majority of recorded points are within the Itatiaia National Park and were overlapping (Fig. 1), indicating it as the area where the species is most commonly sighted.

**Figure 1.**
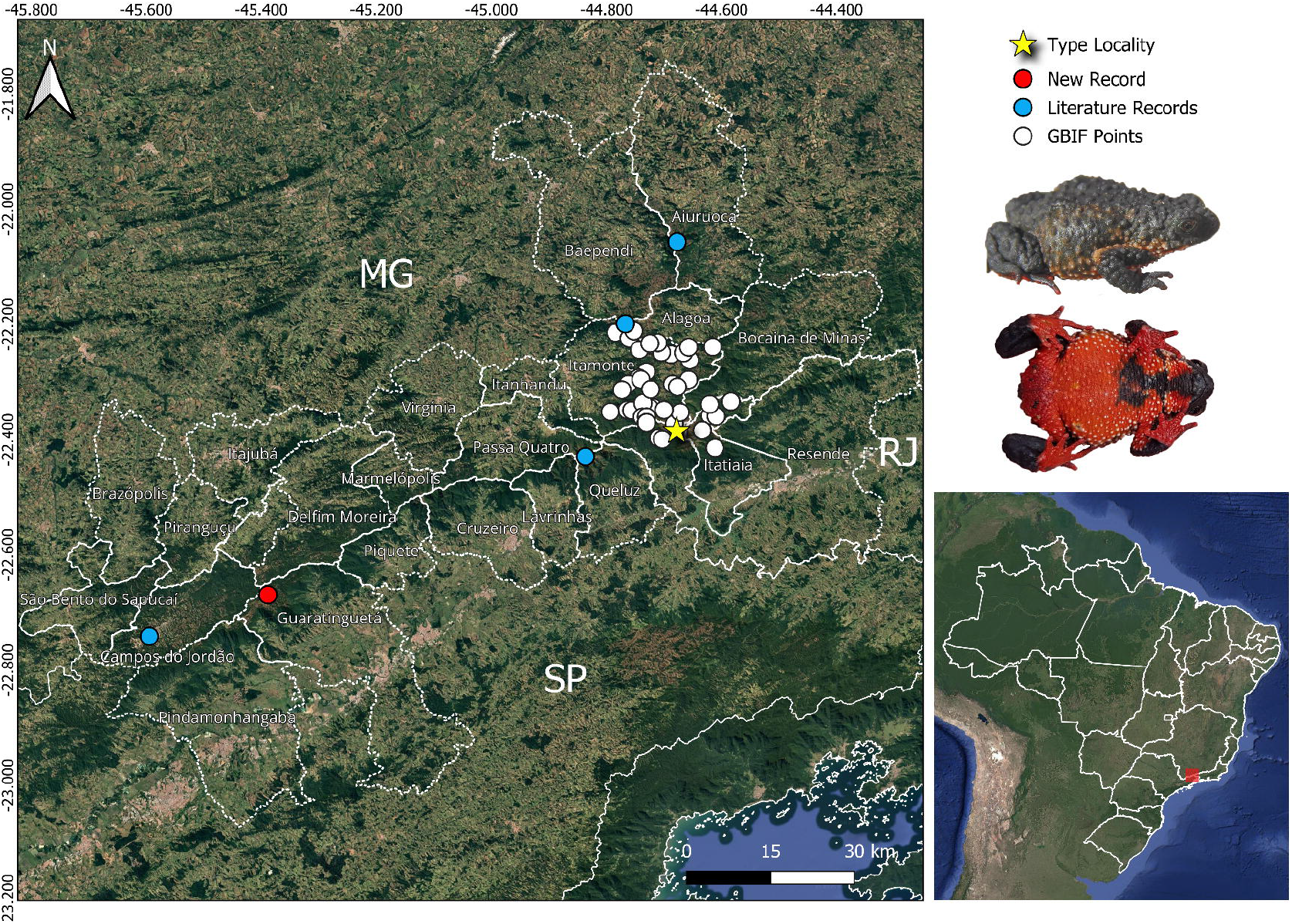
Updated geographic distribution of *Melanophryniscus moreirae* with records from scientific literature and valid records available in the Global Biodiversity Information Facility (GBIF). Solid lines indicate state boundaries, while dashed lines represent municipal boundaries. The municipalities shown correspond to those within the species’ potential range. Abbreviations: MG = Minas Gerais; RJ = Rio de Janeiro; SP = São Paulo.

One possible explanation is that the park is frequently visited by researchers, tourists, and wildlife observers, who are responsible for most of the species records we found.

Until 2006, *M. moreirae* was believed to be endemic to Itatiaia National Park. However, Marques et al. (2006) reported the first record for the state of São Paulo, at Pico da Pedra da Mina in municipality of Queluz (22°25’ S, 44°50’ W, elev. 2,570 m), followed by a record by Weber et al. (2007) in the municipality of Aiuruoca, Minas Gerais (22°03’ S, 44°40’ W, elev. 2,138 m). In 2014, in a study modeling the climatic niche and the effects of climate change on *Melanophryniscus* distribution, Zank et al. (2014) reported a new locality in São Paulo, in the municipality of Campos do Jordão. However, this record did not include collection details such as coordinates, elevation, or specific reference that could be traced back to its origin, making it the only record of the species for Campos do Jordão. Ortiz et al. (2017) added another record in the municipality of Alagoa, Minas Gerais, specifically in Serra do Papagaio National Park (22°11’ S, 44°46’ W, WGS 84, elevation 2,220 m). Despite efforts to obtain additional information, such as searches in databases like *splink*.*net*, review of references, GBIF, supplementary materials, as well as contacting authors who mentioned the point in Campos do Jordão in their distribution, we did not receive responses or details. Given the circumstances, the point in Campos do Jordão (Fig. 1) was recorded using the municipality centroid (22°44’ S, 45°35’ W, WGS 84, elev. 1,598 m), but detailed information is required since it is the only one in the literature without a precise location or associated with any type specimen. For this reason, it is possible that our record in Guaratinguetá (SP) is the southwestern-most record in relation to the type locality and the furthest from the type locality (79 km). However, the region where we found the species borders the municipality of Campos do Jordão, so it is possible that the species also occurs in the municipality, although we could not find evidence to support this record.

In the analysis of the temporal series of *M. moreirae* records, the species was most commonly recorded in November 1964 by different researchers (Fig. 2). Indeed, the species’ reproductive period occurs between September and December, with a peak in October, coinciding with the onset of the rainy season that fills temporary pools used as breeding sites by the species (Van Sluys and Guido-Castro 2011). After this period, it is still possible to occasionally find some individuals until mid-April (pers. comm., Aosf). However, with the arrival of the dry and cold period in the high mountains, with temperatures below zero, individuals enter a state of dormancy (Carvajalino-Fernández et al. 2013). This behavior, characterized by reduced metabolism during low temperatures and consequent decreased locomotion, causes them to bury themselves in burrows, known as hibernacula, at a depth of 5– 15 cm below the ground until the cold and dry period ceases (Carvajalino-Fernández et al. 2013). This behavior is an adaptive mechanism to cope with extreme cold (Segadas-Vianna and Dau 1965; Carvajalino-Fernández et al. 2013). In part, our data are consistent with what is known about the natural history of the species, although we found a decrease in encounters during the rainy season in December. It is possible that GBIF data, when observed on a temporal scale, may carry observation biases that do not necessarily reflect the life history of the species. On the other hand, we did not find records for the driest and coldest time of the year, with only one record in May, which is consistent with the literature (Segadas-Vianna and Dau 1965; Carvajalino-Fernández et al. 2013). We propose that new studies with *M. moreirae* or other species may help evaluate whether historical data and platforms that compile species occurrence data can reflect the reproductive biology of the species.

**Figure 2.**
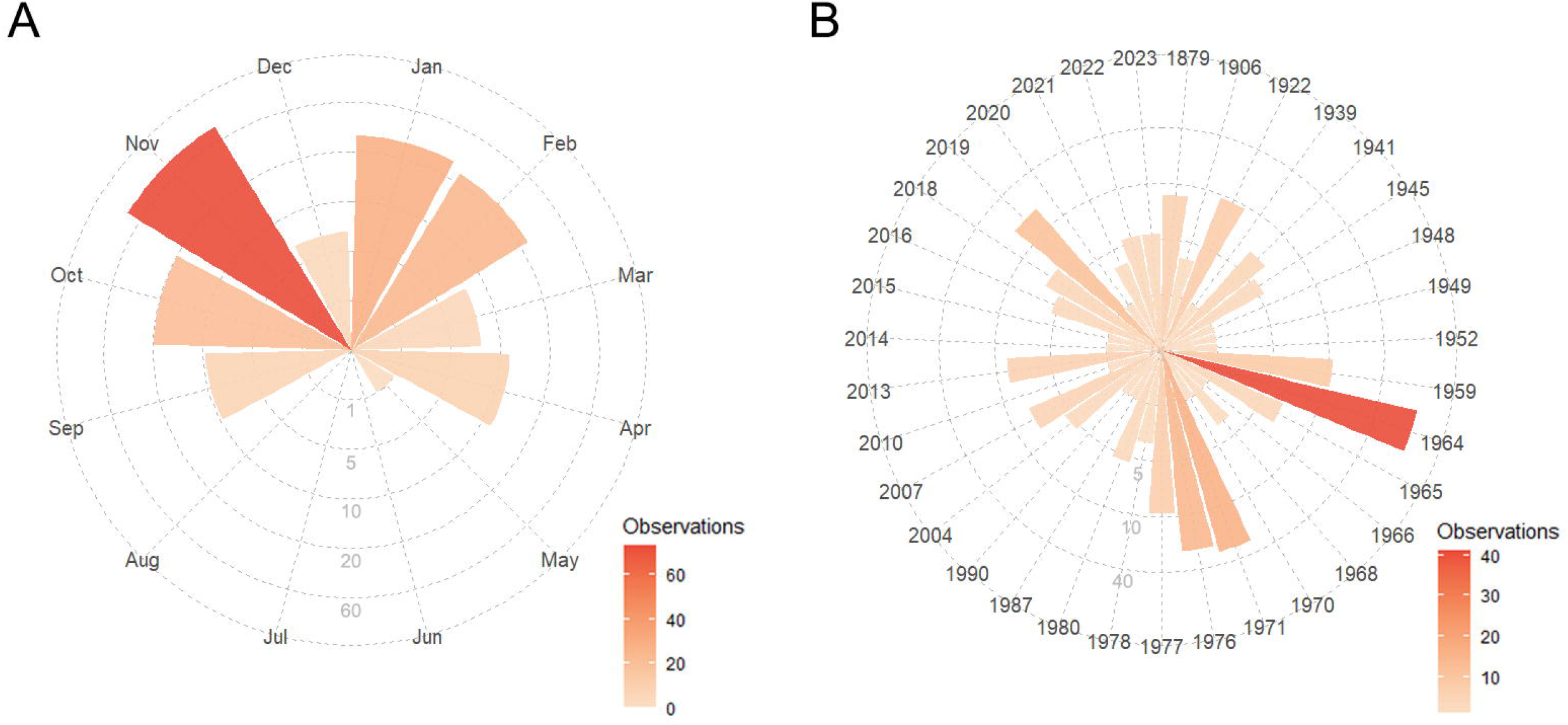
Frequency of records (or observations) of *Melanophryniscus moreirae* obtained from the Global Biodiversity Information Facility (GBIF) platform (A) per month and (B) across years (1879–2023).

Regarding the frequency analysis among elevation classes, it is important to note that although most records are concentrated between 1,800 m and 2,000 m (Fig. 3), the literature establishes a broader average range from 1,800 m to 2,500 m (Guix et al. 1998; IUCN 2023).

**Figure 3.**
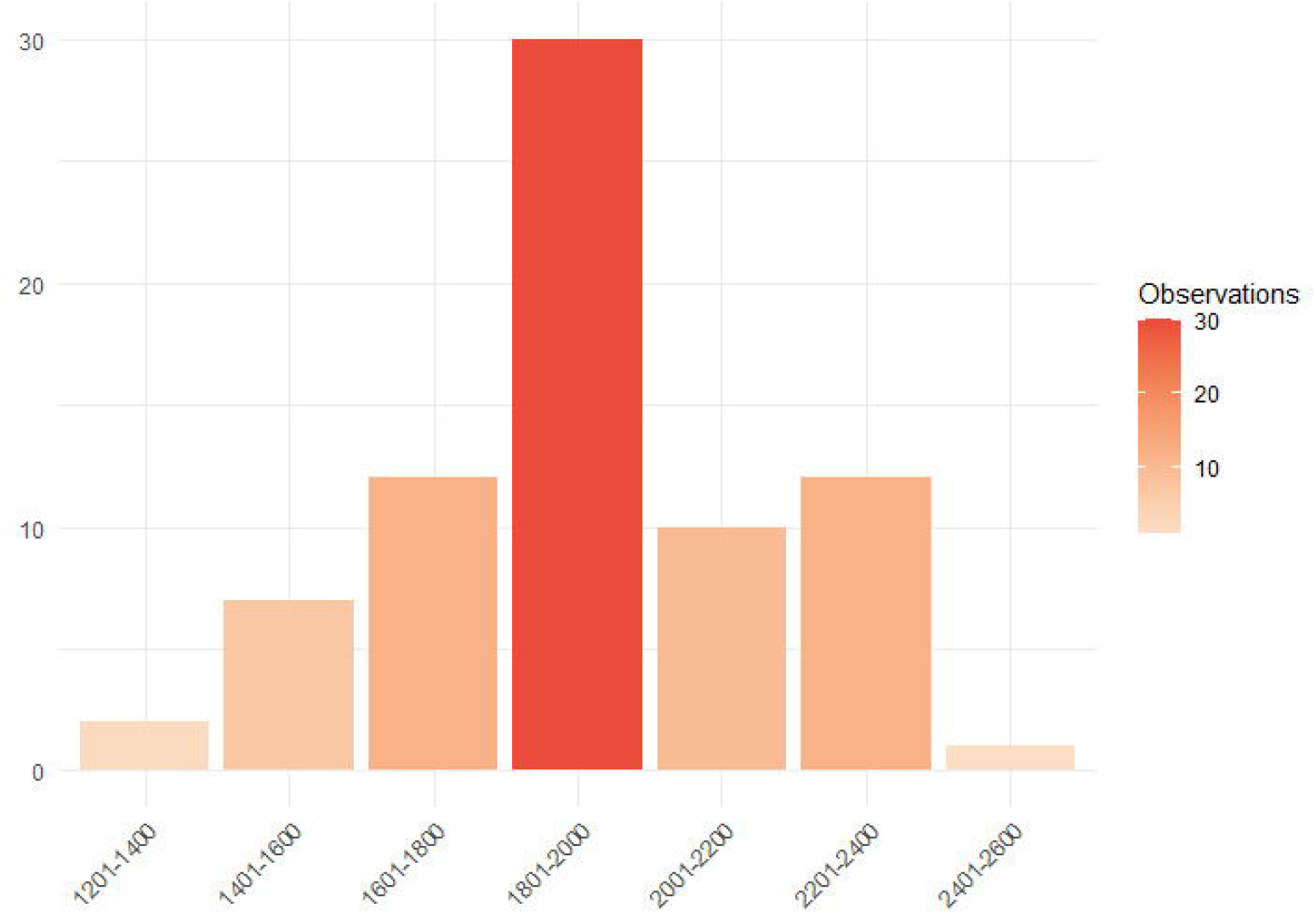
Frequency of *Melanophryniscus moreirae* records between elevational classes based on the coordinates available in the Global Biodiversity Information Facility (GBIF) data.

Additionally, validated records (supplementary material) suggesting the species occurrence at elevations below 1,800 m, possibly starting from 1,200 m, have been found. In this regard, this result may affect the calculations of the species occurrence area. However, it is crucial to emphasize that although these data have been reviewed and confirmed on iNaturalist, there is still some uncertainty regarding the accuracy of the coordinates, as they are sourced from external sources. For this reason, a cautious approach is necessary when interpreting and using these data for further analyses, and more studies should be directed towards evaluating the species abundance along the elevational gradient.

Our study conducted an exhaustive search to include the highest number of records of the “sapo-flamenguinho” a threatened and charismatic species that occurs exclusively in the southern portion of the Serra da Mantiqueira. Until further taxonomic studies incorporating molecular techniques are conducted, there appear to be disjunct populations of *M. moreirae* in locations distant from the Itatiaia plateau, also occurring on the Campos do Jordão plateau. Additionally, we emphasize the importance of including precise data, such as exact occurrence coordinates and, if possible, adding information about elevation, habitat, climate, date, and even time of encounter, especially when dealing with threatened species. The use of citizen science in support of scientific research, if it follows pre-established criteria and quality standards, can contribute to long-term species monitoring. Thus, it could significantly contribute to the conservation and understanding of species’ distributions, as well as to natural history studies. For this reason, it is essential for the scientific community to try to consider these records in their work, as well as to strengthen the capacity of public managers to encourage this practice among visitors and residents near parks and green areas.

## Acknowledgement

We dedicate this study to the memory of our dear friend Pedro Luiz Dixon de Carvalho, owner of “Tao do Gomeral”, who provided us with information about the occurrence of the species in the high-elevation fields of the neighborhood. We would also like to extend our thanks to Erick Siqueira Costa, whose presence and support during the exhausting climb and crossing were fundamental to finding the toad. Finally, we would like to thank Mariana Pedrozo for reviewing the English and making suggestions on the manuscript. MTM is grateful for the fellowship granted by the São Paulo Research Foundation (FAPESP #2023/14506-5).

## Supplementary material

Spreadsheet compiling occurrence data of *Melanophryniscus moreirae* separated into two sheets: literature and Global Biodiversity Information Facility.

